# Lag phase length of IAPP amyloid formation predicts the duration of β-cell toxicity

**DOI:** 10.64898/2026.04.22.720252

**Authors:** Annette Plesner, Ping Cao, Daniel P. Raleigh, Andisheh Abedini

## Abstract

Aggregation of islet amyloid polypeptide (IAPP) contributes to pancreatic β-cell dysfunction in diabetes, yet the identity and temporal persistence of the toxic species remain unresolved. Because amyloid formation proceeds through distinct kinetic phases, relating cytotoxicity to these phases provides a direct strategy to test competing mechanistic models. Here, we combine time-resolved β-cell functional assays with concurrent biophysical measurements to quantitatively link IAPP aggregation kinetics to cellular dysfunction across multiple perturbations and sequence variants. We demonstrate that maximal toxicity occurs during the lag phase and establish aggregation kinetics as a predictive framework for the onset and duration of IAPP proteotoxicity. Across 22 independent experiments spanning more than a 450-fold range in lag times, the duration of β-cell dysfunction scales linearly with lag phase length. Perturbations that alter aggregation kinetics, including changes in concentration, temperature, and sequence, predictably shift both the onset and duration of toxicity. Early lag-phase intermediates of the diabetes-associated S20G-IAPP mutant are more toxic than corresponding species formed by wild-type h-IAPP. Together, these results show that aggregation kinetics quantitatively define the temporal window of IAPP proteotoxicity and support a model in which transient pre-fibrillar intermediates, rather than mature fibrils, dominate β-cell dysfunction.

## INTRODUCTION

Aggregation of islet amyloid polypeptide (IAPP, amylin) contributes to β-cell dysfunction and loss in type 2 diabetes and in islet transplantation failure [1–5]. Although IAPP amyloidosis has long been linked to β-cell stress, inflammation, and apoptosis [6–17], the identity of the toxic species remains unresolved. Competing models have implicated mature fibrils, fibril growth, and soluble pre-amyloid intermediates that are formed during the lag phase or via secondary nucleation in the growth phase as the dominant contributors to toxicity [13, 18–28]. These models make distinct, testable predictions regarding the timing of toxicity relative to aggregation kinetics.

Amyloid formation proceeds through three phenomenological kinetic phases: a lag phase characterized by transient oligomer formation, a growth phase dominated by fibril production and secondary nucleation, and a saturation phase in which fibrils coexist with soluble peptide [1] (**Figure 1A**). If pre-fibrillar intermediates are the principal toxic species, toxicity should be maximal during the lag phase and decline as these species are depleted into fibrils. In contrast, if mature fibrils or fibril growth drive toxicity, no simple quantitative relationship between lag phase duration and cytotoxicity would be expected.

**Figure 1.**
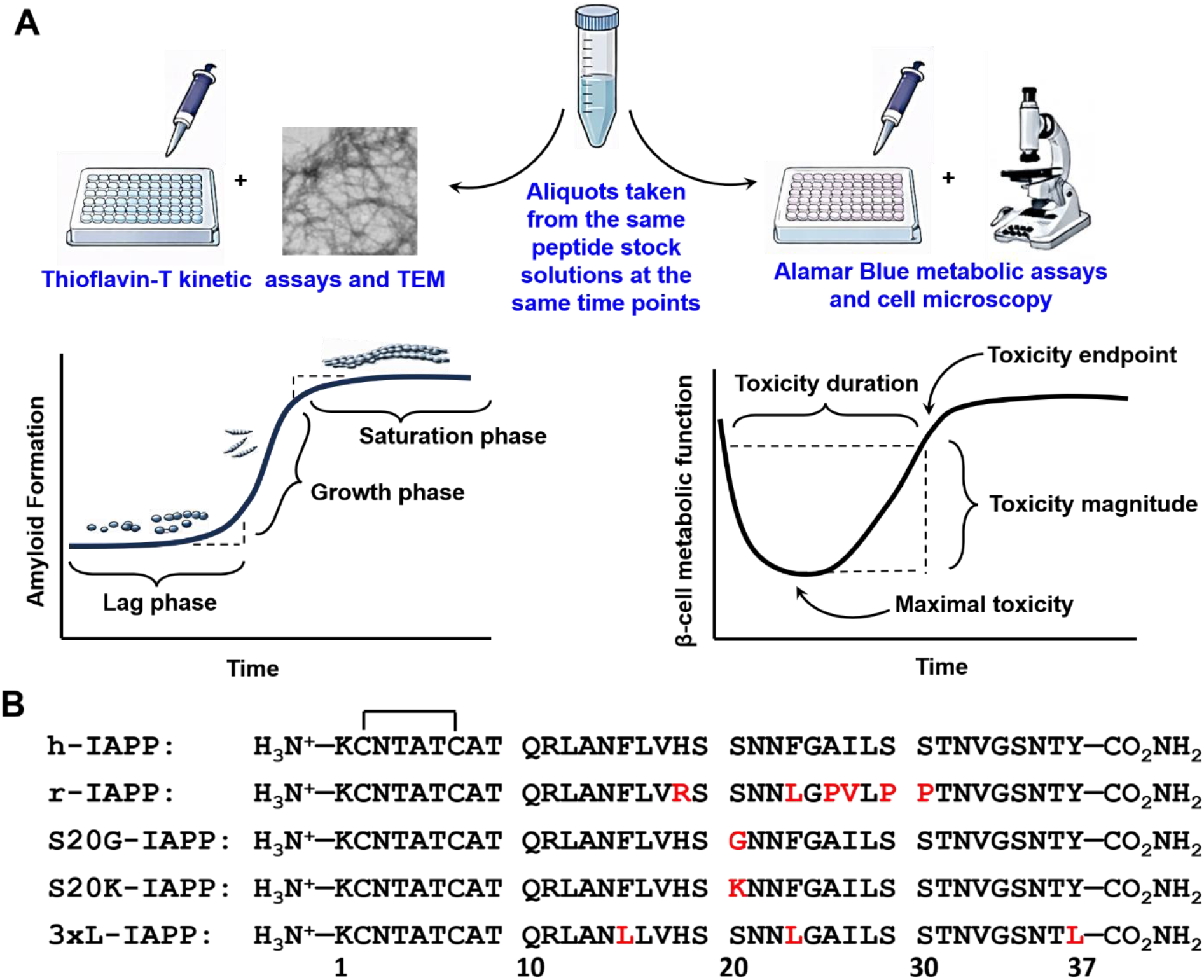
Experimental design and IAPP variants used in the study. **(A)** Schematic of the time-resolved workflow. IAPP peptides were dissolved in 20 mM Tris-HCl, pH 7.4, and incubated at 15 °C or 25 °C. Aliquots were removed at defined times for concurrent thioflavin-T, TEM, Alamar Blue, and light microscopy measurements. Left diagram: The three kinetic phases of amyloid formation. Right diagram: operational definitions of peak toxicity, toxicity duration, and maximal toxicity. **(B)** Primary sequences of h-IAPP, r-IAPP, S20G-IAPP, S20K-IAPP, and 3xL-IAPP. The polypeptides contain a conserved disulfide bond between residues 2 and 7, and an amidated C-terminus. Residues differing from wild-type h-IAPP are highlighted.

Here, we directly test these competing models by combining time-resolved measurements of β-cell dysfunction with concurrent biophysical analysis of IAPP aggregation under conditions that systematically alter lag phase duration. By varying peptide concentration, temperature, and primary sequence, including disease-associated and designed mutants with widely different aggregation kinetics [29–35] (**Figure 1B**), we assess whether lag phase length quantitatively predicts the timing and duration of cytotoxicity. Across all conditions tested, the duration of β-cell dysfunction scales linearly with lag phase length. These findings establish aggregation kinetics as a quantitative framework for predicting the temporal window of IAPP proteotoxicity and support a model in which transient pre-amyloid intermediates are the principal drivers of β-cell dysfunction. While this approach does not directly resolve the molecular identity of the toxic species, it provides a quantitative framework to distinguish between competing mechanistic models.

## RESULTS

To determine whether IAPP-induced β-cell dysfunction is linked to specific stages of aggregation, we systematically conducted 22 independent experiments spanning lag times from less than 1 h to more than 450 h. Time-resolved studies were carried out using previously validated protocols [17, 19]. In each experiment, peptides were dissolved in buffer at physiological pH (pH 7.4) and incubated for varying amounts of time at 15 °C or 25 °C. Aliquots were removed at defined time points over the course of aggregation and analyzed in parallel for amyloid formation (thioflavin-T fluorescence and TEM) and β-cell function (Alamar Blue assays and cell microscopy). This design directly correlates aggregation state to biological outcome, and avoids the ambiguity inherent in endpoint toxicity assays whereby peptide is added immediately after dissolution (time-zero) and aged in the presence of cells. We defined lag time as the time to 10% of the total thioflavin-T intensity change, and toxicity as a reduction in β-cell metabolic function to 80% or less of the response to negative controls. These thresholds were selected on the basis of control experiments with non-amyloidogenic, non-toxic r-IAPP and buffer-treated cells [19] (**Figures 1 and S1**).

### Aggregation kinetics quantitatively define the temporal window of β-cell dysfunction

The rate of h-IAPP amyloid formation is concentration- and temperature-dependent. As expected, lowering the concentration of h-IAPP increased the length of the lag phase, as did lowering the temperature, while increasing the concentration shortened the lag phase. These studies enabled us to examine lag phases ranging from approximately 3 to 25 h under otherwise comparable conditions. In all cases, shortening the lag phase shifted toxicity to earlier time points and compressed its duration, whereas lengthening the lag phase delayed toxicity and prolonged its duration (**Figures 2 and S2**). At the lowest peptide concentration, which displayed the longest lag phase, the onset of toxicity was correspondingly delayed (**Figure 2A and 2B**). A similar trend between lag phase length and toxicity duration was observed when amyloid formation was slowed by lowering the temperature from 25 °C to 15 °C (**Figure S2**). Characterization of samples collected at the time points used for β-cell assays revealed no detectable amyloid fibrils at toxic time points, as judged by thioflavin-T binding assays and TEM (**Figures 2B, 2C and S2B-R**). Our results are consistent with previous studies examining freeze-dried h-IAPP species produced at pH 5.5, 10 °C and 20 μM h-IAPP [22]. In all cases, toxicity consistently occurred at time points before the observation of mature fibrils and preceded the thioflavin-T-defined growth and saturation phases. Maximal toxicity was observed during the lag phase rather than during the later growth or saturation phases. These results temporally associate β-cell dysfunction with pre-fibrillar intermediates, and establish a direct relationship between lag phase length and the onset and duration of toxicity.

**Figure 2.**
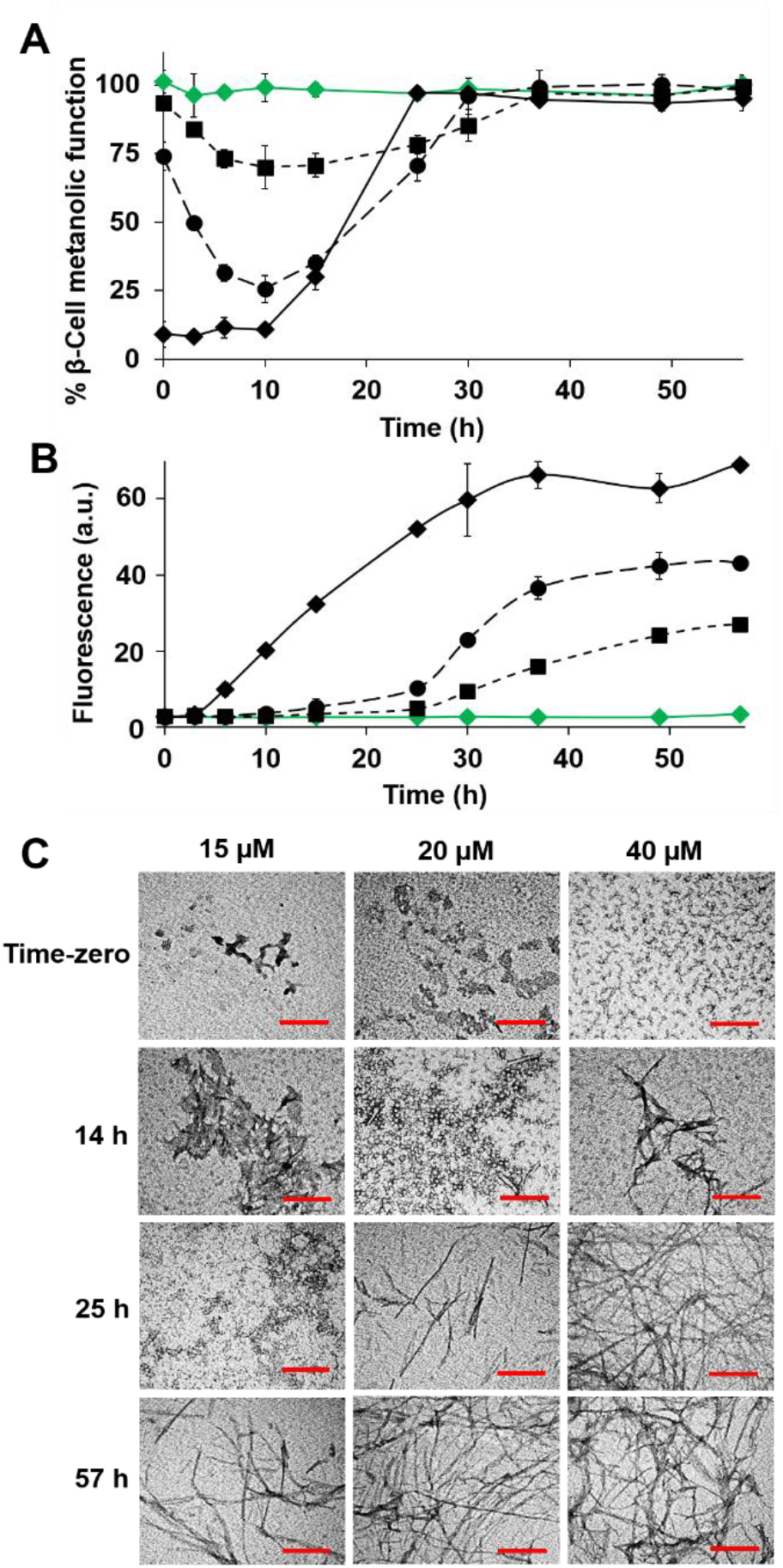
Concentration-dependent changes in the kinetics of h-IAPP amyloid formation correlate with changes in the time course of toxicity. Time-resolved dose-response studies monitoring: **(A)** loss in β-cell metabolic function after treatment of INS-1 β-cells with h-IAPP species produced during the time course of amyloid formation, and **(B)** amyloid formation kinetics at 25 °C as measured by thioflavin-T binding assays. Samples contain 15 μM h-IAPP (black ◼/--- -), 20 μM h-IAPP (black ●/- - -), 40 μM h-IAPP (black ♦/—) and 40 μM r-IAPP (negative control, green 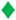/—). The same color and symbol codes are used in all panels. Final peptide concentrations after dilution into β-cell assays were 10.5, 14, and 28 μM, respectively. In panels A and B, the data represent means ± SEM from three to six replicate wells per condition and at least three independent experiments. Some of the error bars are the same size or smaller than the symbols in the graphs. **(C)** TEM data collected at different time points during the time course of amyloid formation by the different peptide solutions used in studies shown in panels A and B (Scale bars: 200 nm). Data presented in panels A-C were collected at the same time points using aliquots from the same amyloid formation assays at 25 °C.

### IAPP sequence variants compress or extend the toxicity window in proportion to lag phase duration

To further extend the range of our experiments, we analyzed IAPP sequence variants with well-characterized effects on aggregation. The rationally designed mutants, which either decrease or increase the rate of amyloid formation compared to h-IAPP, extended the accessible kinetic range to more than 450-fold in lag phase duration (**Figures 3, S3 and S4**). Mutations at position 20 are known to modulate the kinetics of h-IAPP amyloid formation [29], and a naturally occurring Ser20Gly missense variant in the human IAPP gene is associated with increased risk for diabetes [4, 29–33]. Substitution of Ser-20 with Gly (S20G-IAPP) largely abolishes the detectable lag phase, while substitution with a Lys (S20K-IAPP) increases the lag phase length by a factor of more than 30 relative to h-IAPP at the same concentration (**Figures 3 and S3**) [29]. Relative to h-IAPP, faster-aggregating S20G-IAPP produced an earlier onset and shorter duration of transient toxicity, whereas slower-aggregating S20K-IAPP delayed toxicity onset and peak toxicity and extended the toxicity window (**Figures 3, S3 and S4**). Notably, maximal toxicity differed between variants, indicating that the magnitude and duration of toxicity are separable properties. The maximal cytotoxicity induced by the bulk population of early pre-amyloid S20G-IAPP species is significantly greater than that produced by h-IAPP at the same concentration (**Figure S5**), consistent with prior observations in cultured islets that S20G-IAPP is more toxic than h-IAPP [31]. The greater maximal toxicity of S20G-IAPP could reflect either a larger population of toxic intermediates, higher intrinsic toxicity of those intermediates, or both. We do not distinguish between these possibilities here. Although the increased amyloidogenicity of S20G-IAPP could be interpreted as evidence for fibril-mediated toxicity, our time-resolved data argue against that model by showing that toxicity is maximal before fibril growth and mature amyloid fibril accumulation.

**Figure 3.**
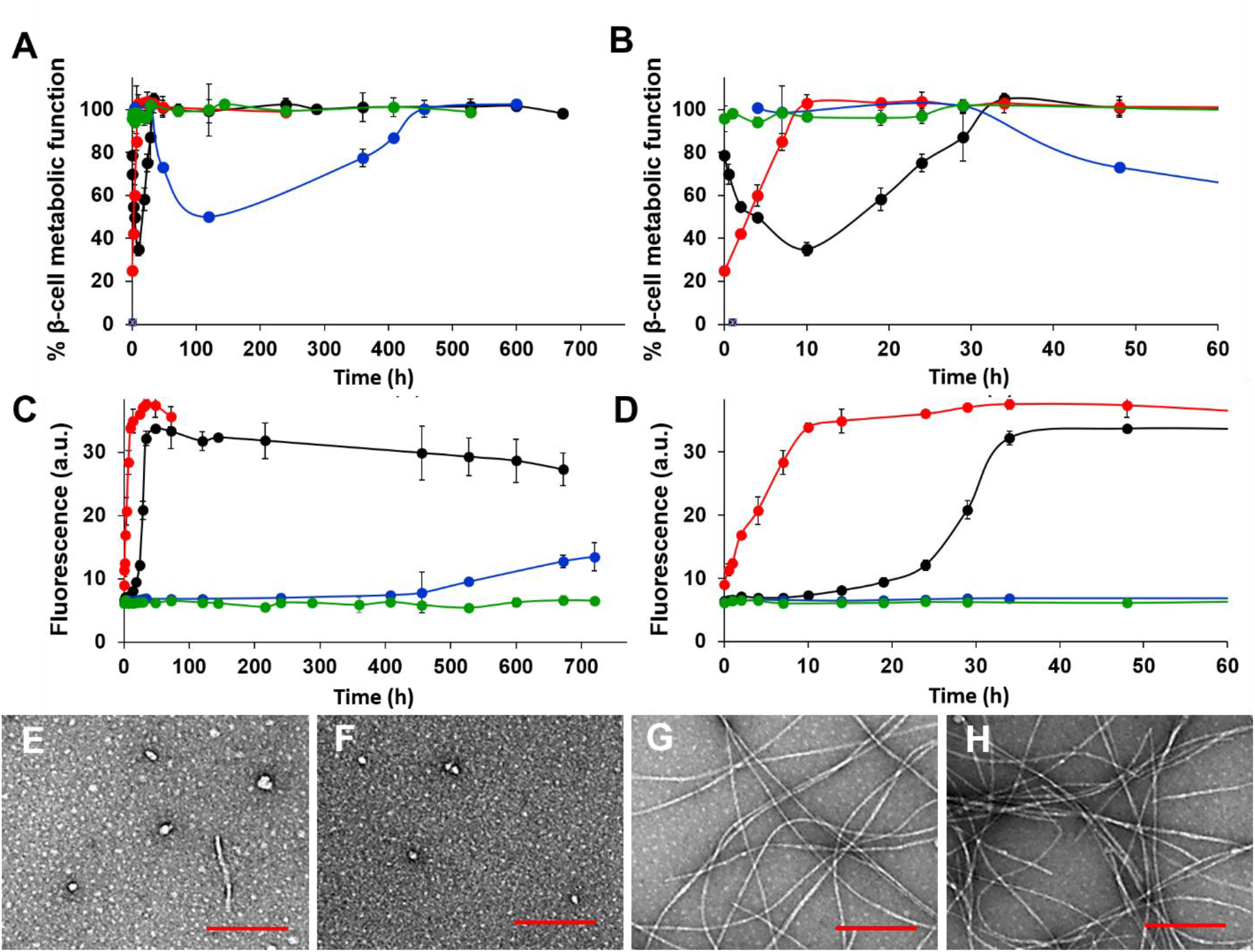
S20 point mutants of h-IAPP reveal a direct relationship between the length of the lag phase of amyloid formation and the onset and duration of cytotoxicity. **(A)** Time-resolved Alamar Blue metabolic assays of β-cells treated with different kinetic species produced during the time course of amyloid formation: WT h-IAPP (black ●), S20G-IAPP (red 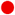), S20K-IAPP (blue 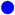) and r-IAPP (negative control, green 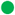). The final peptide concentration after diluting peptide stock aliquots into β-cell assays was 14 μM. **(B)** The first 60 min of the time-resolved Alamar Blue metabolic data shown in panel A. **(C)** Thioflavin-T monitored kinetics of amyloid formation at 25 °C carried out concurrently with Alamar Blue assay shown in panel A using aliquots from the same peptide stock solutions. **(D)** The first 60 min of the time-resolved thioflavin-T kinetics data shown in panel C. The same color and symbol codes are used in all panels. Some of the error bars in panels A-D are the same size or smaller than the symbols in the graphs. Data represent means ± SEM from three to six replicate wells per condition and at least three independent experiments. **(E-H)** TEM image of **(E)** S20G-IAPP (time-zero), **(F)** toxic S20K-IAPP (mid lag-phase), **(G)** non-toxic S20G-hIAPP fibrils, and **(H)** non-toxic S20K-IAPP fibrils (Scale bars: 200 nm). TEM data were collected using aliquots from the same amyloid formation assays which were used in experiments shown in panels A-D.

We also compared the behaviors of h-IAPP and S20K-IAPP to a previously reported triple mutant of h-IAPP in which Phe-15, Phe-23 and Tyr-37 were substituted to Leu (3xL-IAPP) [19, 34, 35]. 3xL-IAPP forms amyloid more slowly than h-IAPP but more rapidly than S20K-IAPP. The lag phase of 3xL-IAPP is 22-fold longer than that of h-IAPP and 1.6-fold shorter than S20K-IAPP, at the same concentration (**Figure S3 and S4**) [19, 34, 35]. Our studies show that the increase in the lag phase length of 3xL-IAPP relative to h-IAPP is accompanied by an increase in the time required to reach the maximum level of toxicity, and a significant increase in the total duration of toxicity. However, the maximum toxicity achieved at any point by 3xL-IAPP is significantly less than the maximum value achieved for h-IAPP or S20K-IAPP (**Figure S5**). This difference could reflect either a smaller population of toxic oligomers formed by 3xL-IAPP than by h-IAPP or S20K-IAPP, lower intrinsic toxicity of those oligomers, or both. These possibilities cannot be distinguished with the current experimental design, as no non-perturbing quantitative assay exists to measure the population of transient toxic oligomers in cell culture. Interestingly, the final thioflavin-T intensity for the 3xL-IAPP mutant was similar to that of the S20K-IAPP mutant at the same concentration, which was lower than that for h-IAPP or S20G-IAPP. It is well established that the final thioflavin-T intensity in an amyloid assay is not a quantitative measure of fibril mass, and the data presented here further show that endpoint thioflavin-T intensity does not predict the onset, duration, or magnitude of proteotoxicity. These findings indicate that sequence-dependent factors influence both the population and intrinsic properties of toxic intermediates, independent of their lifetime.

### Lag phase duration quantitatively predicts the toxicity window

Analysis of all 22 independent experiments revealed a very strong linear relationship between lag phase duration and the duration of β-cell toxicity (*R*^*2*^ = 0.92, *P* < 3.6 × 10^-12^) across a kinetic range spanning less than 1 h to more than 450 h (**Figure 4 and Table S1**). This relationship remained statistically significant when the analysis was restricted to experiments with lag times of 25 h or less, indicating that the correlation is not driven by the longest-timescale data points. Lag phase duration also predicted both the endpoint of toxicity and the time of peak β-cell dysfunction. A strong linear relationship was observed between lag phase duration and the time at which toxicity ended (*R*^*2*^ = 0.94, *P* < 3.6 × 10^-14^), as well as between lag-phase T50 and the time of peak toxicity (*R*^*2*^ = 0.65, *P* < 5.8 × 10^-6^) (**Figures 5 and S6**). A nonparametric Spearman rank analysis yielded the same overall conclusion, confirming a strong monotonic relationship between lag phase duration and toxicity duration (*ρ* = 0.81, *P* < 3.24 × 10^-6^), including when the analysis was restricted to data points with lag times of 25 h or less (*ρ* = 0.71, *P* < 1.4 × 10^-4^). Together, these analyses indicate that lag phase duration predicts the onset, peak, and termination of β-cell dysfunction. These data demonstrate that aggregation kinetics do not merely correlate qualitatively with toxicity, but quantitatively predict the temporal window during which toxic species are present.

**Figure 4.**
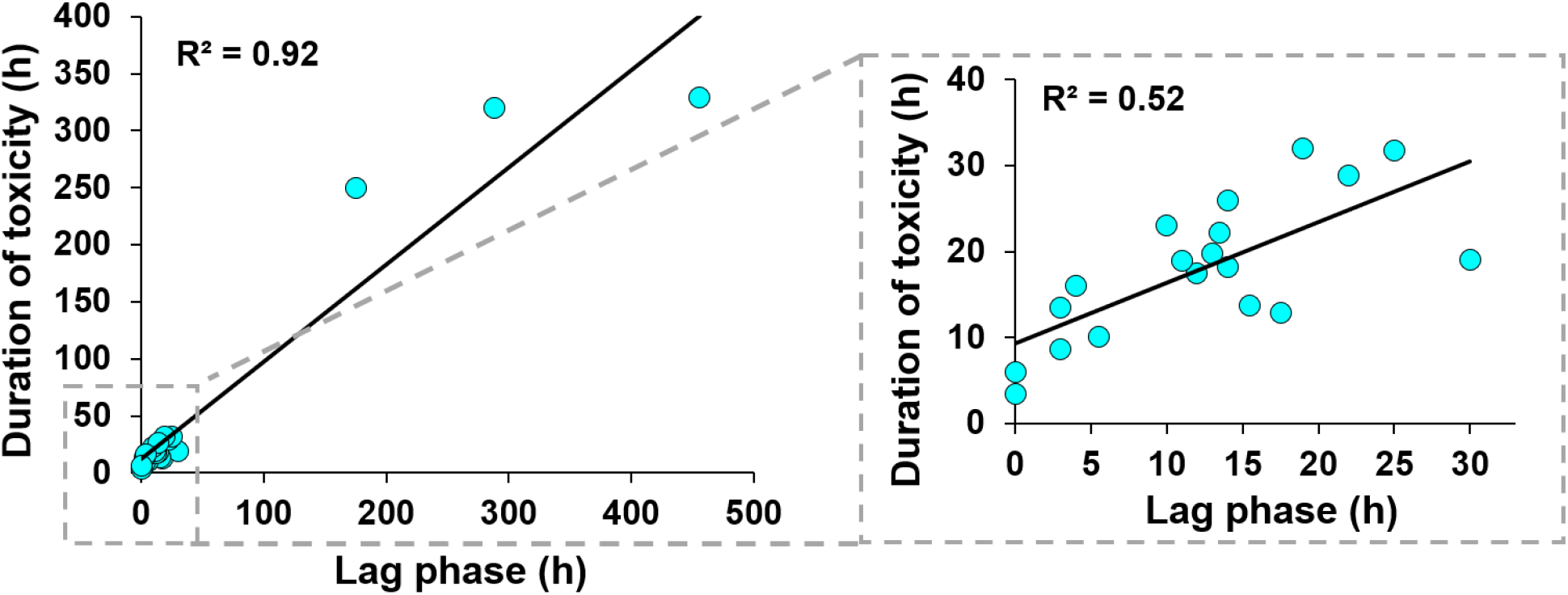
Lag phase duration quantitatively predicts the duration of β-cell toxicity. Linear regression analysis of concurrent thioflavin-T binding assays and Alamar Blue metabolic assays were carried out for the 22 independent experiments, including those presented in Figures 2, 3, S2, S3, S4 and Table S1. The measured lag phase lengths spanned a range from <1 h to >450 h. Data demonstrate a highly statistically significant linear relationship between the length of the lag phase and the duration of toxicity (*R*^*2*^ = 0.92, *P* < 3.6 × 10^-12^). Inset shows the data at early time points. The linear relationship for data collected at early time points is also highly statistically significant (*R*^*2*^ = 0.52, *P* < 1.4 × 10^-4^).

**Figure 5.**
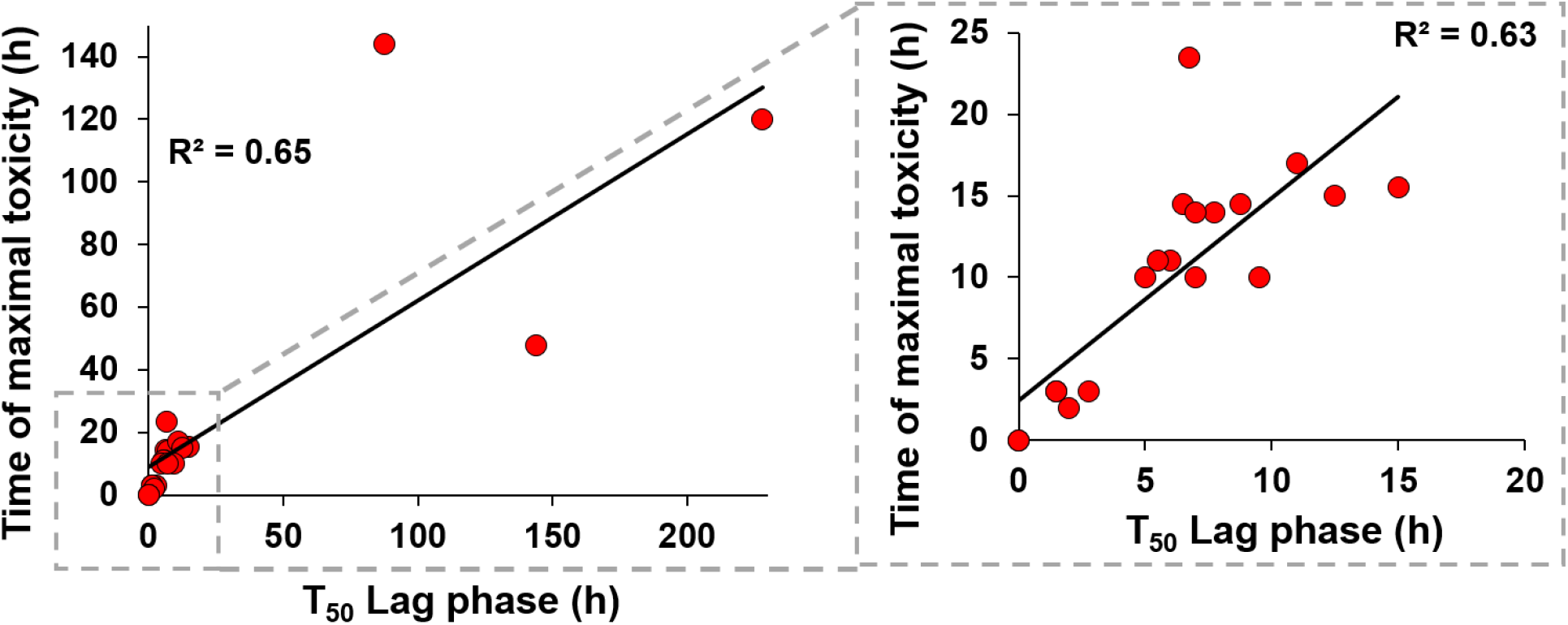
Lag phase T_50_ predicts the time of maximal β-cell toxicity. Linear regression analysis of concurrent thioflavin-T binding assays and Alamar Blue metabolic assays were carried out for the 22 independent experiments, including those presented in Figures 2, 3, S2, S3, S4, and Table S1. The measured lag phase lengths spanned a range from <1 h to >450 h. Data indicate a highly statistically significant linear relationship between the T_50_ of the lag phase length and the time point of maximal toxicity (*R*^*2*^ = 0.65, *P* < 5.1 × 10^-6^). The linear relationship for data collected at early time points, shown in the inset, is also highly statistically significant (*R*^*2*^ = 0.63, *P* < 1.1 × 10^-5^).

## DISCUSSION

Our central finding is that the lag phase of IAPP amyloid formation quantitatively defines the temporal window of β-cell dysfunction. This relationship holds across multiple perturbations, including concentration, temperature, and sequence variation, spanning well over two orders of magnitude in aggregation timescale.

These data support a model in which transient pre-fibrillar intermediates are the principal mediators of β-cell dysfunction. Toxicity arises and peaks during the lag phase, and declines in the growth phase as toxic intermediates are depleted into fibrils. Across all conditions, maximal toxicity occurred before the midpoint of the growth phase. Notably, for slowly aggregating variants such as S20K-IAPP and 3xL-IAPP, toxicity declined before the onset of the growth phase, further arguing against a dominant contribution from fibril growth under these conditions. The decrease in detectable toxicity during the growth phase and its absence in the saturation phase are consistent with a model in which fibril formation reduces toxicity by sequestering harmful intermediates.

Importantly, our results distinguish between the duration and magnitude of toxicity. The S20G-IAPP variant, which aggregates rapidly and exhibits little to no detectable lag phase, produces higher peak toxicity over a shorter time window, consistent with previous observations that this mutation enhances amyloidogenicity and cytotoxicity [29–33]. In contrast, slower-aggregating variants such as S20K-IAPP and 3xL-IAPP extend the lag phase and the duration of β-cell dysfunction, but differ in their maximal effects on β-cell function [19, 34, 35]. These observations indicate that sequence-dependent factors modulate the population and intrinsic toxicity of oligomeric intermediates, independent of their temporal persistence.

These data establish a temporal relationship between aggregation kinetics and toxicity but do not directly define the structural identity of the toxic intermediates. Whether the toxic species characterized here are “on-pathway” to fibril formation remains unresolved. Prior real-time 2D IR studies suggest that the lag time of h-IAPP amyloid formation is controlled by refolding of a lag-phase oligomer containing transient partial β-sheet structure in the F_23_GAIL_27_ region. Although those studies were performed at higher peptide concentrations than those used here, they raise the possibility that this oligomer is related to the toxic species observed in our assays [36–39]. Our conclusions do not depend on whether these species are on- or off-pathway intermediates. Rather, the data establish that toxicity is temporally linked to pre-fibrillar species rather than fibrils, that maximal toxicity occurs during the lag phase, and that lag phase duration quantitatively predicts the window of toxicity.

More broadly, these findings identify aggregation kinetics as a predictive determinant of the onset and duration of proteotoxicity in nucleation-dependent amyloid systems. While these studies are performed *in vitro*, the identification of a defined kinetic window of toxicity suggests that similar temporal dynamics may govern β-cell exposure to toxic intermediates *in vivo*. Because lag phases are a general feature of amyloid assembly, this principle may extend beyond IAPP to other disease-associated proteins in which transient pre-fibrillar intermediates are implicated as toxic species.

Therapeutically, these results argue for targeting early aggregation intermediates and the kinetic processes that govern their formation and persistence. Interventions that shorten the lifetime or reduce the population of pre-amyloid intermediates will decrease the window of β-cell exposure to toxic species. Conversely, strategies focused solely on mature fibrils are unlikely to reduce toxicity unless they also reduce the population or lifetime of pre-fibrillar species. Thus, lag phase duration is not simply a descriptive feature of amyloid formation; it is a quantitative determinant of proteotoxic exposure.

Despite establishing a quantitative relationship between aggregation kinetics and β-cell dysfunction, several key questions remain. Most notably, the molecular identity and structural features of the toxic pre-fibrillar intermediates are still unclear. Determining whether these species represent a discrete oligomeric ensemble or a continuum of transient conformations, and whether they are obligate on-pathway intermediates or off-pathway species, will be critical for advancing mechanistic understanding. Equally important is the development of non-perturbing quantitative methods to directly measure the population and lifetime of these intermediates in complex biological environments, including living cells and native islet tissue. Another major challenge is translating these in vitro kinetic relationships to the *in vivo* setting, where local peptide concentration, membrane composition, and cellular stress responses may modulate both aggregation pathways and toxicity. Finally, integrating aggregation kinetics with downstream cellular signaling pathways, including inflammatory and stress-response networks, will be essential to connect molecular events to β-cell failure in disease. Resolving these questions will sharpen the mechanistic basis of IAPP proteotoxicity and help define therapeutic strategies that selectively target the earliest, most pathogenic stages of amyloid formation.

## ACKNOWLEDGEMENTS

We thank Professor Christopher Newgard at Duke University for generously providing rat INS-1 β-cells and Dr. Daeun Noh for assistance with data analysis.

## MATERIALS AND METHODS

### Peptide synthesis and purification

Wild-type human IAPP (h-IAPP), rat IAPP (r-IAPP), and IAPP variants (S20G-IAPP, S20K-IAPP, and 3xL-IAPP) were prepared by solid-phase peptide synthesis using standard 9-fluorenylmethoxycarbonyl (Fmoc) chemistry on a 0.10 mmol scale with a CEM Liberty microwave peptide synthesizer, as previously described [19, 40, 41]. Peptides were synthesized on 5-(4′-Fmoc-aminomethyl-3′,5-dimethoxyphenol) valeric acid (PAL-PEG) resin to yield a C-terminal amide. Following cleavage and deprotection, crude peptides were purified by reverse-phase high-performance liquid chromatography (HPLC). Peptide identity and purity were confirmed by analytical HPLC and mass spectrometry, as described previously [19, 40, 41]. All peptides contained the native disulfide bond between residues 2 and 7 and were used without further modification.

### Preparation of peptide solutions

Lyophilized peptides were dissolved in 20 mM Tris-HCl buffer (pH 7.4) immediately prior to use. Solutions were handled in a manner designed to minimize pre-aggregation. Aggregation reactions were initiated by dilution into buffer at the indicated concentrations and incubated under quiescent conditions at either 15 °C or 25 °C. To directly relate aggregation state to biological activity, aliquots were removed from the same peptide solutions at defined time points throughout the aggregation time course and used immediately for biophysical and cellular assays. Aggregation experiments were conducted at either 15 °C or 25 °C, as specified in each experiment. Peptide concentration was varied to modulate aggregation kinetics. Following dilution into cell culture medium, final peptide concentrations in β-cell assays ranged from 10.5 to 56 μM.

### Thioflavin-T fluorescence (ThT) assays

Amyloid formation was monitored using ThT fluorescence assays as previously described [19]. ThT was added to peptide samples, and fluorescence was measured over time to track amyloid formation kinetics. Lag time was defined operationally as the time required to reach 10% of the total ThT fluorescence change. This definition is consistent with prior work and enables quantitative comparison across conditions [19].

### Transmission electron microscopy (TEM)

Samples collected at defined time points during aggregation were analyzed by transmission electron microscopy to assess fibril formation. Specimens were prepared and imaged using established protocols [19]. TEM analysis was performed in parallel with ThT measurements using aliquots from the same peptide solutions, enabling direct correlation between fibril formation and aggregation kinetics.

### Cell culture

Immortalized rat INS-1 β-cells were used for all cell-based assays and maintained under standard culture conditions as previously described [19]. Cultures were routinely monitored for Mycoplasma contamination, and only Mycoplasma-free cells were used in experiments. Cell identity was verified using standard institutional cell-line authentication procedures.

### β-cell viability and metabolic function assays

β-cell metabolic function was assessed using Alamar Blue reduction assays, following previously established protocols [19]. Cells were exposed to peptide samples for 5 h prior to measurement of metabolic activity. Peptide samples were obtained from stock solutions at defined time points during the aggregation process. Importantly, peptide samples were pre-aggregated *in vitro* prior to addition to cells, and aliquots corresponding to specific aggregation states were applied to β-cells. This design avoids ambiguity associated with time-zero addition experiments in which aggregation occurs in the presence of cells. Toxicity was defined as a reduction in β-cell metabolic function to ≤80% of the response observed in buffer-treated controls. This threshold was established based on control experiments with non-amyloidogenic r-IAPP and untreated cells [19].

### Light microscopy

Cell morphology was assessed by light microscopy following peptide treatment, using previously described methods [19]. Imaging was performed in parallel with metabolic assays to qualitatively evaluate cellular integrity and morphology.

### Experimental design and replication

All experiments were performed using time-resolved sampling of the same aggregation reactions, enabling direct coupling of aggregation state to biological outcome. Each condition included 3–6 technical replicates per experiment, and all experiments were repeated independently at least three times. Data are presented as mean ± SEM unless otherwise indicated.

### Data analysis

Lag phase duration, time to toxicity onset, time of peak toxicity, and duration of toxicity were extracted from time-resolved ThT and Alamar Blue datasets. Linear regression and nonparametric analyses were performed as described in the Results. Statistical significance was assessed using the tests indicated in the relevant figure legends and supplemental figures. Regression analyses were performed on untransformed data; results were qualitatively unchanged upon log transformation of lag time (data not shown).

### Data availability

All data generated or analyzed during this study are included in the manuscript, supporting information, and source data files.

### Funding Sources

This work was supported by grants: 17SDG33410350 (A.A.) from the American Heart Association (AHA); GM078114 (D.P.R.) from the National Institutes of Health (NIH).

