## Supplementary material for "Lag phase length of IAPP amyloid formation predicts the duration of β-cell toxicity": Supp Info: Abedini et al_Supp Info_BioRxiv_042226.pdf

**SUPPORTING FIGURES**

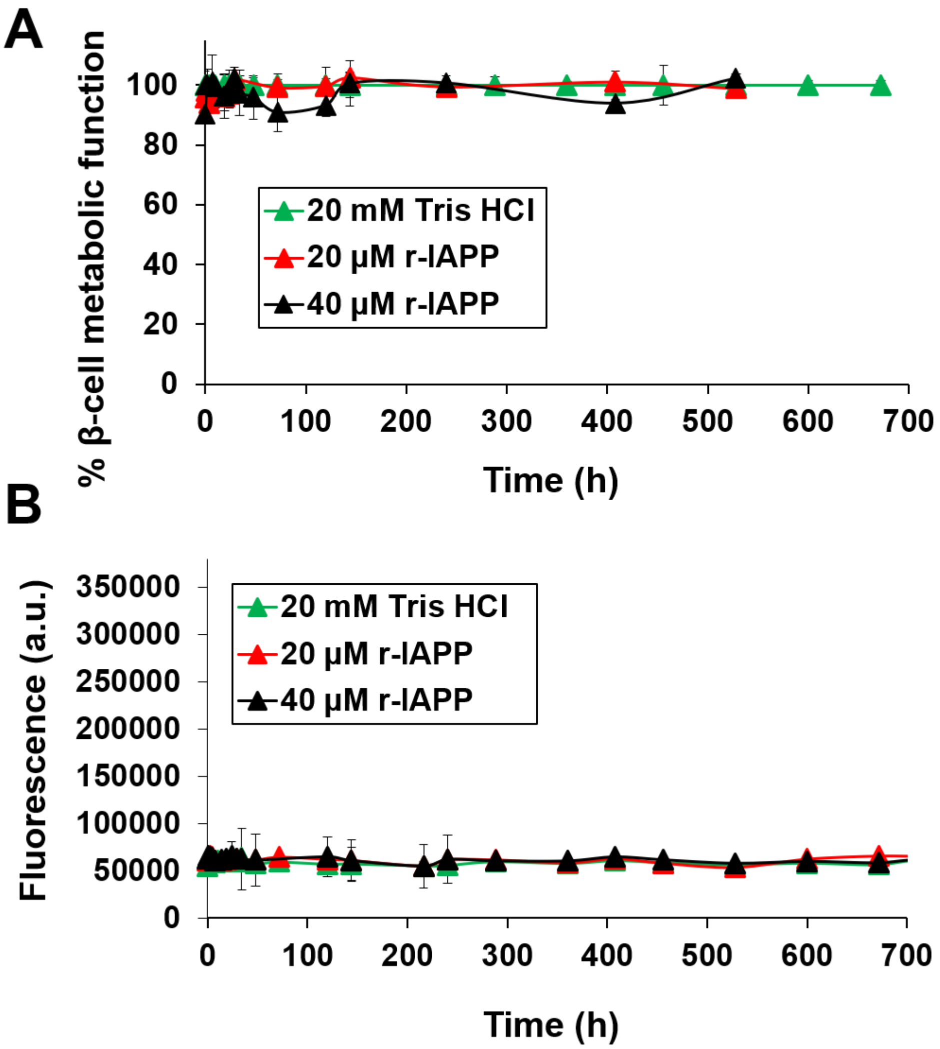


**Figure S1. Time-resolved dose-response studies of toxicity and amyloid formation by rat IAPP (r-IAPP). (A)** Alamar Blue metabolic assays measure the response of β-cells after 5 h treatment with aliquots of peptide or buffer control solutions. **(B)** Thioflavin-T fluorescence (unnormalized)-monitored kinetics of amyloid formation carried out concurrently with metabolic assays shown in panel A. Some of the error bars in panels A and B are the same size or smaller than the symbols in the graphs. Data represent means ± SEM of three to six replicate wells per condition and a minimum of three replicate experiments per group.

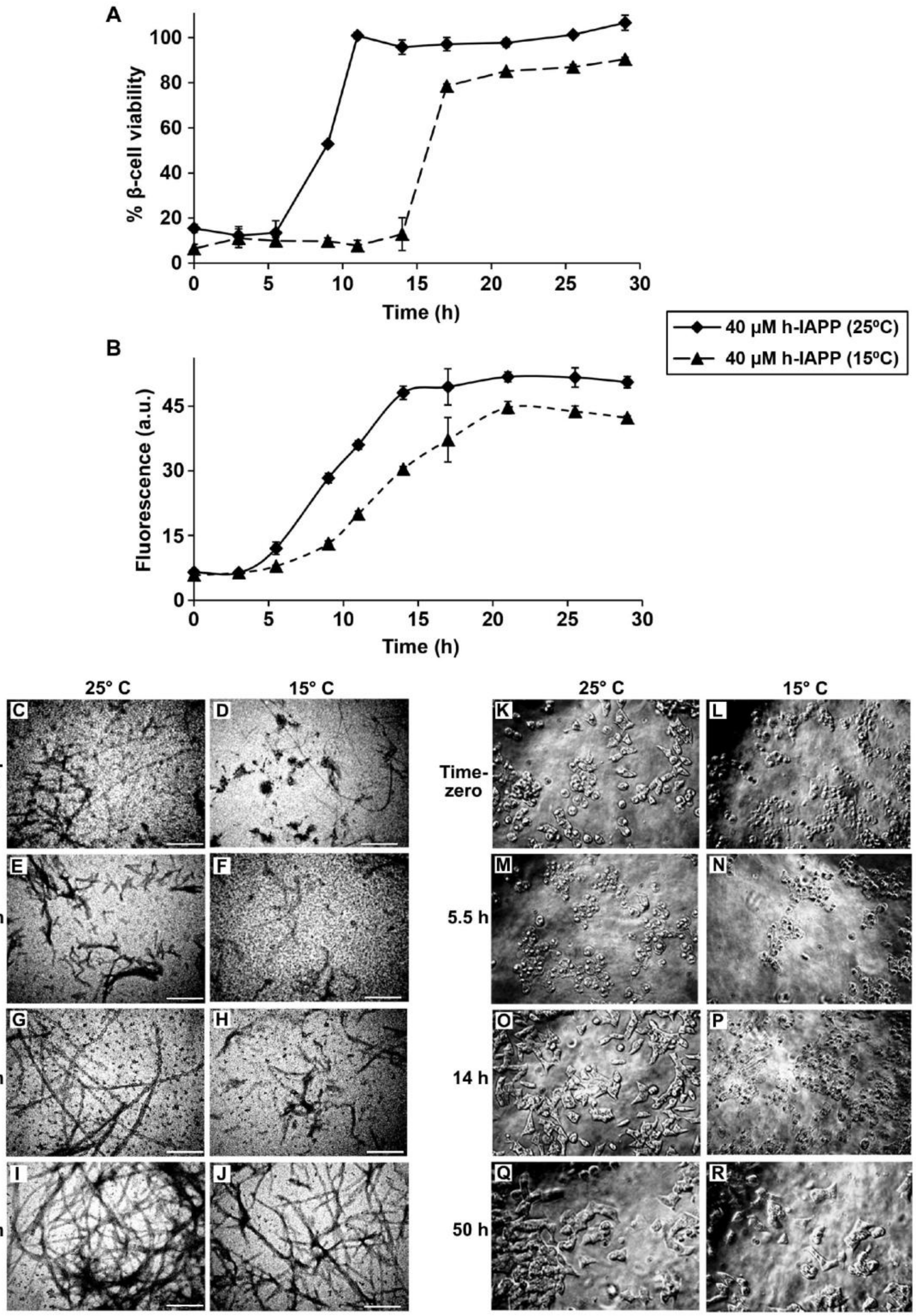


**Figure S2. Changes in the kinetics of h-IAPP amyloid formation induced by a change in temperature lead to correlated changes in the time course of toxicity.** Time-resolved studies using: **(A)** Alamar Blue metabolic assays to monitor β-cell viability and, **(B)** thioflavin-T binding assays to monitor amyloid formation kinetics. Peptide solutions contained 40 μM peptide and were incubated at either 25 °C (black ♦/—) or 15 °C (black ▲/----). The same color and symbol codes are used in all panels. Final peptide concentration after dilution of peptide solutions into β-cell assays was 28 μM. Some of the error bars in panels A and B are the same size or smaller than the symbols in the graphs. Although the proportionality between lag phase length and toxicity duration is less exact at 15 °C than at 25 °C, lowering the temperature consistently lengthened the lag phase and prolonged the toxicity window. Data represent means ± SEM of three to six replicate wells per condition and three replicate experiments per group. **(C-J)** TEM data collected at the same time points as those assessed in panels A and B: **(C)** Time-zero, 25 °C, **(D)** Time-zero, 15 °C, **(E)** 5.5 h, 25 °C, **(F)** 5.5 h, 15 °C, **(G)** 14 h, 25 °C, **(H)** 14 h, 15 °C, **(I)** 50 h, 25 °C and, **(J)** 50 h, 15 °C (Scale bars: 200 nm). **(K-R)** Light microscopy images of β-cells after treatment with the different h-IAPP kinetic species shown in panels C-J: **(K)** Time-zero, 25 °C, **(L)** Time-zero, 15 °C, **(M)** 5.5 h, 25 °C, **(N)** 5.5 h, 15 °C, **(O)** 14 h, 25 °C, **(P)** 14 h, 15 °C, **(Q)** 50 h, 25 °C and **(R)** 50 h, 15 °C. Data presented in panels A-R were collected at the same time points in concurrent experiments using aliquots from the same respective amyloid formation assays containing 40 μM peptide.

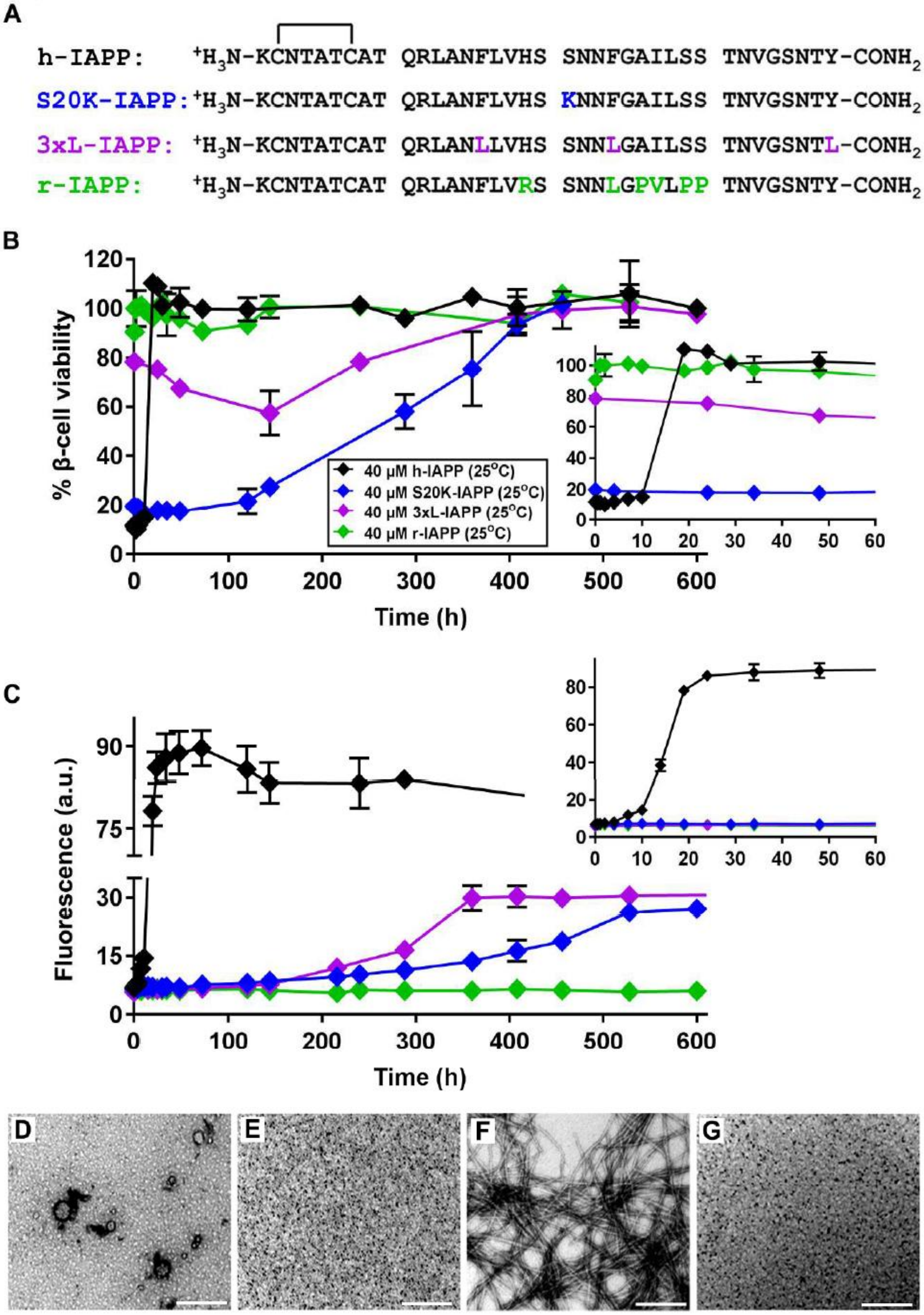


**Figure S3. The kinetics of amyloid formation and toxicity by wild type and variants of h-IAPP are directly linked. (A)** Primary sequences of h-IAPP, S20K-IAPP, 3xL-IAPP and r-IAPP. Amino acid positions that differ from WT h-IAPP are indicated by color. **(B)** Time-resolved Alamar Blue metabolic assays showing β-cell viability after treatment with different kinetic species produced during amyloid formation: h-IAPP (black ♦), 3xL-IAPP (purple ♦), S20K-IAPP (blue ♦) and r-IAPP (green ♦). **(C)** Thioflavin-T monitored kinetics of amyloid formation by h-IAPP, 3xL-IAPP, S20K-IAPP and r-IAPP at 25 °C. The same color and symbol codes are used in all panels. Data represent means ± SEM of three to six replicate wells per condition and three replicate experiments per group. Insets in panels B and C show the data at early time points. Some of the error bars in panels B and C are the same size or smaller than the symbols in the graphs. **(D-G)** TEM data showing: **(D)** toxic, mid-lag phase species of 3xL-IAPP; **(E)** r-IAPP at time points corresponding to the mid-lag phase of 3xL-IAPP; **(F)** 3xL-IAPP amyloid fibrils (saturation phase); and **(G)** r-IAPP at time points corresponding to the saturation phase of 3xL-IAPP (Scale bars: 200 nm). Alamar Blue metabolic assays, thioflavin-T binding assays and TEM studies were carried out at the same time using aliquots from the same respective amyloid formation assays carried out at 25 °C, containing 40 μM peptide. Final peptide concentration after dilution of peptide solutions into β-cell assays was 28 μM. Note that kinetic data and TEM images for 3xL-IAPP presented in panels B and C are also shown on a different scale in our previously published article by Abedini et al. and are included here for direct comparison across variants [19].

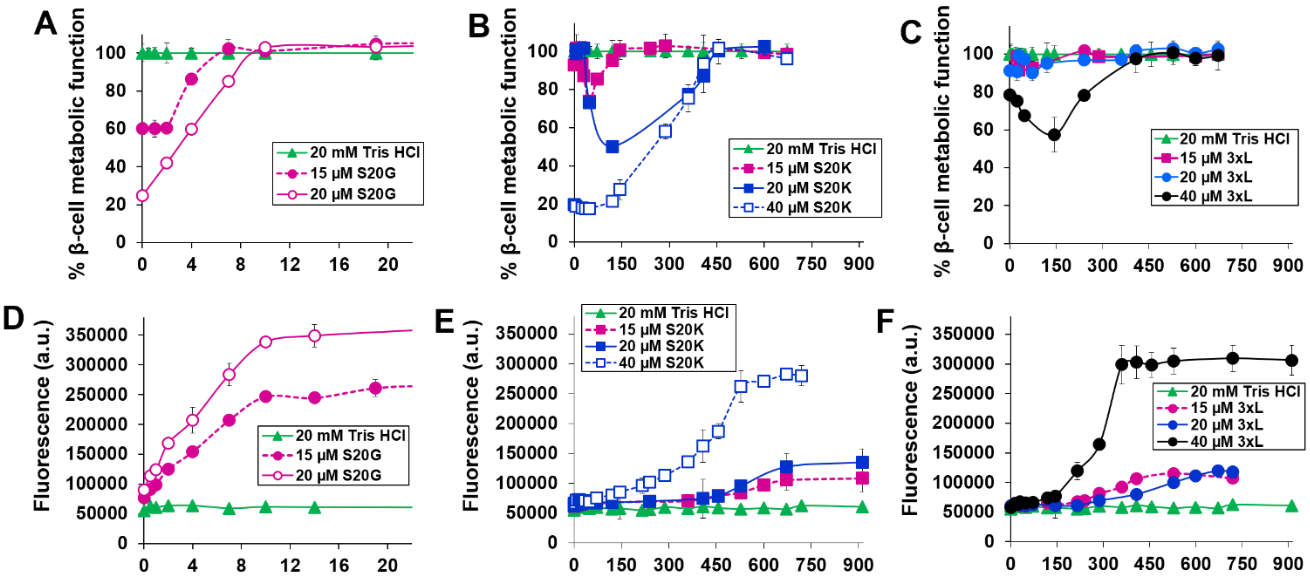


**Figure S4. Time-resolved dose-response studies of toxicity and amyloid formation by S20G-IAPP, S20K-IAPP and 3xL-IAPP. (A-C)** Alamar Blue metabolic assays measure the response of β-cells after 5 h treatment with aliquots of peptide or buffer control (pH 7.4) solutions taken at different times over the course of amyloid formation at 25 °C. **(D-F)** Thioflavin-T fluorescence (unnormalized)-monitored kinetics of amyloid formation carried out concurrently with metabolic assays shown in panels A-C. The final peptide concentrations on cells after dilution of the peptide stocks into cell culture were 10.5 µM (from 15 µM samples), 14 µM (from 20 µM samples), and 24 µM (from 40 µM samples). Data represent means ± SEM of three to six replicate wells per condition and three replicate experiments per group. For some data points, error bars are smaller than the size of the symbols in the graphs. Data for 40 µM 3xL-IAPP were previously reported in Abedini et al. and are included here for direct comparison across variants [19].

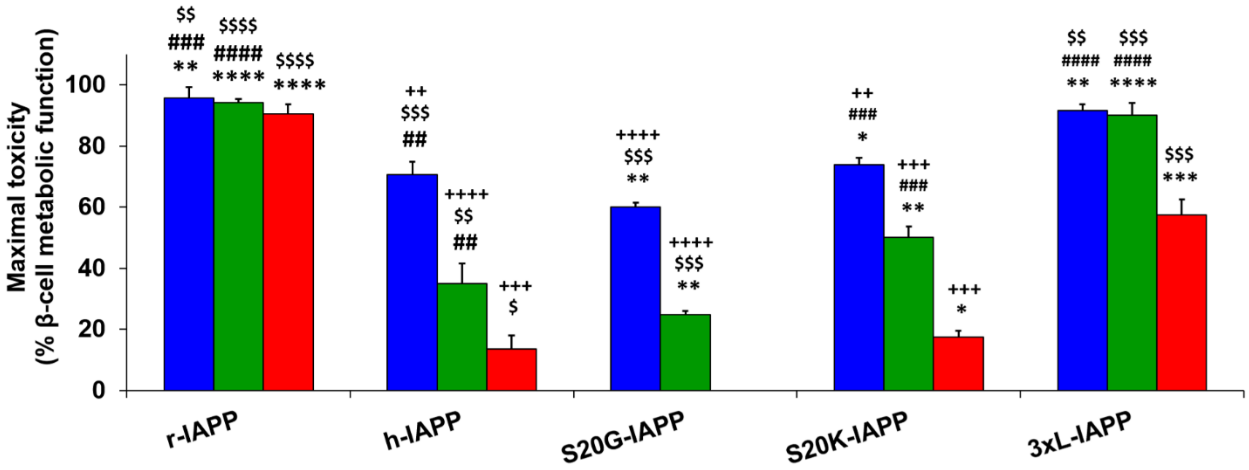


**Figure S5. Comparison of the maximal level of toxicity elicited by r-IAPP, h-IAPP, S20G-IAPP, S20K-IAPP, and 3xL-IAPP over the course of amyloid formation.** Bars are color-coded by final peptide concentration on cells: 10.5 μM (blue), 14 μM (green), and 24 μM (red). The time-resolved data from which averages were obtained for each peptide condition are presented in Figures 2, 3, S2, S3, S4 and Table S1. Data is normalized to buffer-treated cells and represent means ± SEM of three to six replicate wells per condition and a minimum of three replicate experiments per group. Statistical significance was determined using a two-tailed Student's t-test. *$P < 0.05$, **$P < 0.01$, ***$P < 0.001$, ****$P < 0.0001$ versus h-IAPP at the same concentration. ##$P < 0.01$, ###$P < 0.001$, ####$P < 0.0001$ versus S20G-IAPP at the same concentration. \$$P < 0.05$, \$\$$P < 0.01$, \$\$\$$P < 0.001$, \$\$\$\$$P < 0.0001$ versus S20K-IAPP at the same concentration. ++$P < 0.01$, +++$P < 0.001$, ++++$P < 0.0001$ versus 3xL-IAPP at the same concentration.

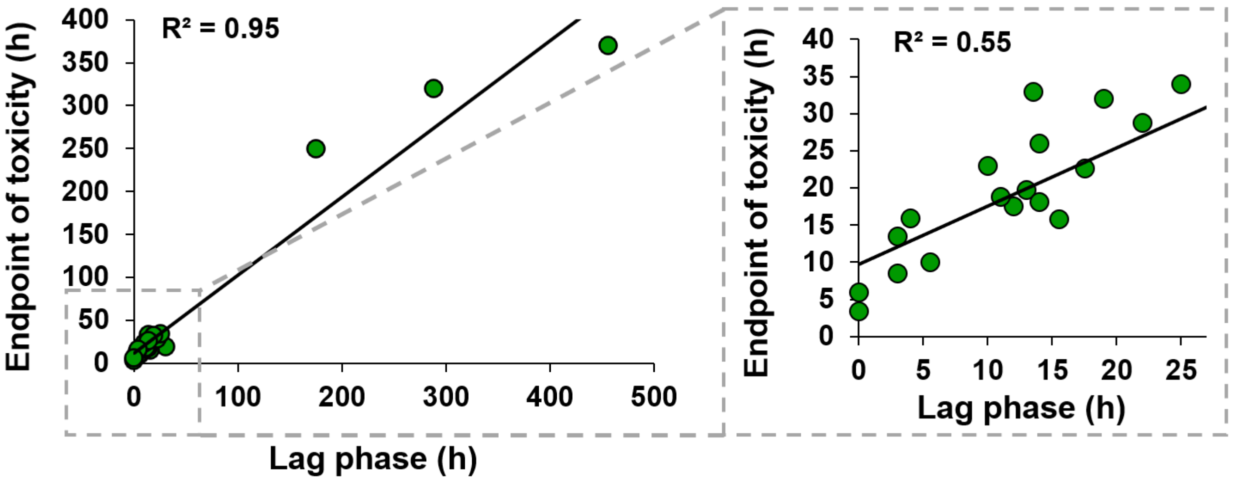


**Figure S6. Conditions that alter the rate of h-IAPP amyloid formation show a direct and statistically significant correlation between the length of the lag phase and the endpoint of toxicity.** Linear regression of a total of 22 independent measurements involving different conditions and different variants of h-IAPP, including those presented in Figures 2, 3, S2, S3, S4 and Table S1, was used to study the relationship between the kinetics of amyloid formation and cytotoxicity. The measured lag phase lengths spanned a range from <1 h to >450 h. Data demonstrate a highly statistically-significant linear relationship between the length of the lag phase and the time point at which the duration of toxicity ends ($R^2 = 0.95$, $P < 3.6 \times 10^{-14}$). The linear relationship for data collected at early time points, shown in the inset, is also highly statistically significant ($R^2 = 0.55$, $P < 8.37 \times 10^{-5}$). Inset shows the data at early time points. The final range of peptide concentrations after dilution of peptide stock solutions into β-cell assays for correlation studies was 10.5 to 56 μM.

| Sample Conditions | Lag phase length (h) | T50 Lag phase length (h) | Time of toxicity onset (h) | Max Toxicity (% metabolic function) | Time of Max Toxicity (h) | Toxicity duration (h) | Toxicity endpoint (h) |
|---|---|---|---|---|---|---|---|
| 15 µM h-IAPP (25C) | 25.0 | 12.5 | 2.2 | 70.7 | 15.0 | 31.8 | 34.0 |
| 20 µM h-IAPP (25C) | 19.0 | 9.5 | 0.1 | 25.6 | 10.0 | 32.0 | 32.0 |
| 20 µM h-IAPP (25C) | 17.5 | 8.8 | 9.8 | 22.1 | 14.5 | 12.9 | 22.7 |
| 20 µM h-IAPP (25C) | 14.0 | 7.0 | 0.0 | 35.0 | 10.0 | 26.0 | 26.0 |
| 20 µM h-IAPP (25C) | 15.5 | 7.8 | 2.2 | 52.9 | 14.0 | 13.7 | 15.9 |
| 30 µM h-IAPP (25C) | 14.0 | 7.0 | 0.0 | 8.4 | 14.0 | 18.2 | 18.2 |
| 30 µM h-IAPP (25C) | 10.0 | 5.0 | 0.0 | 6.4 | 10.0 | 23.0 | 23.0 |
| 40 µM h-IAPP (25C) | 3.0 | 1.5 | 0.0 | 8.3 | 3.0 | 13.5 | 13.5 |
| 40 µM h-IAPP (25C) | 13.0 | 6.5 | 0.0 | 6.3 | 14.5 | 19.8 | 19.8 |
| 40 µM h-IAPP (25C) | 13.5 | 6.8 | 10.8 | 34.6 | 23.5 | 22.2 | 33.0 |
| 40 µM h-IAPP (25C) | 4.0 | 2.0 | 0.0 | 10.1 | 2.0 | 16.0 | 16.0 |
| 40 µM h-IAPP (25C) | 5.5 | 2.8 | 0.0 | 12.3 | 3.0 | 10.1 | 10.1 |
| 80 µM h-IAPP (25C) | 3.0 | 1.5 | 0.0 | 5.7 | 3.0 | 8.6 | 8.6 |
| 20 µM h-IAPP (15C) | 22.0 | 11.0 | 0.0 | 20.4 | 17.0 | 28.8 | 28.8 |
| 40 µM h-IAPP (15C) | 30.0 | 15.0 | 0.0 | 24.6 | 15.5 | 19.0 | 19.0 |
| 40 µM h-IAPP (15C) | 12.0 | 6.0 | 0.0 | 7.8 | 11.0 | 17.5 | 17.5 |
| 15 µM S20G-IAPP (25C) | 0.0 | 0.0 | 0.0 | 60.1 | 0.0 | 3.5 | 3.5 |
| 20 µM S20G-IAPP (25C) | 0.0 | 0.0 | 0.0 | 24.8 | 0.0 | 6.0 | 6.0 |
| 20 µM S20K-IAPP (25C) | 456.0 | 228.0 | 41.0 | 50.1 | 120.0 | 329.0 | 370.0 |
| 40 µM S20K-IAPP (25C) | 288.0 | 144.0 | 0.0 | 17.6 | 48.0 | 320.0 | 320.0 |
| 40 µM 3xL-IAPP (25C) | 175.0 | 87.5 | 0.0 | 57.5 | 144.0 | 250.0 | 250.0 |
| 40 µM h-IAPP / 40 µM I26P-IAPP mixture (25C) | 11.0 | 5.5 | 0.0 | 19.4 | 11.0 | 18.9 | 18.9 |

**Table S1. Summary of 22 independent experiments linking the lag phase of IAPP amyloid formation to β-cell toxicity.** Table of experimental conditions and quantitative parameters extracted from concurrent thioflavin-T amyloid formation assays and Alamar Blue β-cell metabolic assays for wild-type and variant IAPP peptides. Conditions include peptide identity, concentration, temperature, lag phase length, duration of toxicity, time of toxicity onset, level and time of maximal toxicity, and time of end of toxicity in time-resolved assays. These data form the basis for the correlation analyses shown in Figures 4, 5, and S6, demonstrating that the duration of toxicity scales with the length of the amyloid lag phase.
